# Genetically targeted 3D visualisation of *Drosophila* neurons under Electron Microscopy and X-Ray Microscopy using miniSOG

**DOI:** 10.1101/070755

**Authors:** Julian Ng, Alyssa Browning, Lorenz Lechner, Masako Terada, Gillian Howard, Gregory S. X. E. Jefferis

## Abstract

Large dimension, high-resolution imaging is important for neural circuit visualisation as neurons have both long- and short-range patterns: from axons and dendrites to the numerous synapses at their endings. Electron Microscopy (EM) is the favoured approach for synaptic resolution imaging but how such structures can be segmented from high-density images within large volume datasets remains challenging.

Fluorescent probes are widely used to localise synapses, identify cell-types and in tracing studies. The equivalent EM approach would benefit visualising such labelled structures from within sub-cellular, cellular, tissue and neuroanatomical contexts.

Here we developed genetically-encoded, electron-dense markers using miniSOG. We demonstrate their ability in 1) labelling cellular sub-compartments of genetically-targeted neurons, 2) generating contrast under different EM modalities, and 3) segmenting labelled structures from EM volumes using computer-assisted strategies. We also tested non-destructive X-ray imaging on whole *Drosophila* brains to evaluate contrast staining. This enables us to target specific regions for EM volume acquisition.

## Introduction

The ability to identify individual neurons based on their molecular, physiological and connective properties is a fundamental goal in neuroscience. A major part of this effort comes from imaging neurons, from the tracing of intricately branched axons and dendrites that span 10-100s of microns, to the identification of thin neurite endings and synaptic sites that are 10-100s of nanometre in diameter. Identifying all the patterns a single neuron makes reveals not only where synaptic connections reside and what the partner neurons may be but it also gauges its contribution to the functional network it resides in. This requires a volume-imaging approach that not only encompasses millimetre-micron wide dimensions but that also features the appropriate nanometre-scale resolutions. From a functional perspective, it is also crucial that particular neuron types can be identified, based on specific molecular, morphological or physiological characteristics. In this respect, given the complex connectivity patterns within a 3D brain environment, the specific identification and tracing of individual highly interconnected neurons from heterogenous populations of neurites remains a highly challenging task [1].

Many light microscopic options are available to explore the composition and organisation of the brain [2–9]. In many cases, investigators choose probes based on fluorescent proteins (FPs), tracer dyes or dye-attached antibodies, proteins or nucleotides to label neurons and synapses. Subsequent probe localisations can determine each neuron's physiological, gene expression, projection and synaptic patterns. However, with the diffraction limit inherent in light microscopy, resolving densely innervated neurites and their synaptic connectivity patterns is difficult. Strategies such as the GFP reconstitution across synaptic partners (GRASP) technique [10, 11] can help to some extent. With recent super-resolution microscopy, many neural features, such as sub-synaptic structures and resident molecules can be resolved to within 10s of nanometre [12–14]. However, how this will be implemented in densely labelled samples is unclear.

Recent EM techniques are well suited for large volume and high-resolution imaging, capable of capturing every axon and dendrite, synaptic and sub-synaptic feature [15–17]. Various volume EM techniques are currently available [15, 17–20]. Serial sectioning followed by transmission electron microscopy (ssTEM) was the key method used to map the *C. eleg*ans nervous system, and was subsequently used in *Drosophila* and mouse [21–23]. Obtaining large numbers of manually generated, serial aligned and registered images from 100– 1000s of sections is technical demanding, generally involves specialised equipment and requires careful image acquisition and pre-processing procedures [24–29]. Two recent imaging techniques, using serial block face scanning electron microscopy (SBF-SEM or SBEM) and focus ion beam scanning electron microscopy (FIBSEM), offer good alternatives as tissue blocks undergo automated cycles of image acquisition and sectioning [17]. As images are sequentially acquired from the block surface, they are typically well aligned.

One recurrent challenge lies in image segmentation (here also termed as reconstruction) [15, 30]. Dense reconstruction strategies involve tracing all neurites and synaptic features from the image volume, followed by feature and identity annotations [23, 31–33]. Alternatively, sparse mapping strategies reduce the analysis to fewer neurons of interest, by choosing the most relevant features from the image volume for further analysis [22, 34, 35]. In both cases, manual tasks are time-consuming and error-prone, and tracing needs to be repeated and expert-supervised to generate a consensus [15, 36–38]. Novel computational approaches are also being deployed [39–44]. Nonetheless, these methods also have substantial error rates that preclude fully automated global solutions [15]. In practice, while no single procedure is perfect, multiple approaches can work in a complementary manner [15, 31, 37, 43, 45].

EM-visible protein tags that generate electron-dense contrast can help in this effort. Acting as guides, molecularly defined neurons, synaptic sites or targets of choice can be highlighted to map circuits of interest. Such probes include enzyme peroxidases (such as HRP or APEX) [46–49] and photo-sensitiser proteins (such as miniSOG)[50], and photosensitive dyes that associate with proteins [51], or short peptides, such as ReASH [52]. These provide genetic solutions where cells can be made to express these probes *in situ*, which are then processed for EM contrast. To date, while there have been some success, further improvements must be made to extend their targeting capabilities, labelling properties and contrast detection on the neuronal ultrastructure [48, 50, 53–55].

Given the existing challenges in visualising the nervous system, we explored ways by which genetically encoded electron-dense probes can be used to visualise labelled neurons under various volume EM modalities. As EM projects are typically time and labour-intensive we investigated the conditions for optimal labelling, contrast staining, and sample preparation of fly brains. This was also motivated from personal experience: we find that image quality tends to be variable in terms of ultrastructure and contrast staining. Selecting specimens for EM imaging is somewhat error-prone. Without prior knowledge, sub-optimal samples are often used, resulting in wasted efforts. We investigated micro-Computed Tomography (micro-CT, also known as X-ray microscopy or XRM) as a means to assess specimen quality. While biological micro-CT/XRM is typically associated with low contrast detection in soft tissues, heavy metal and osmium-based contrast can be detected, deriving useful cell biological and neuroanatomical information [56–60]. With recent improvements and the availability of laboratory-based scanners, biological materials can be routinely scanned, yielding sub-micron resolutions at large dimensions (0.3-10 millimetre, depending on magnification)[59, 60]. Knowing which area to acquire EM images from is paramount as the field dimension is typically restricted, especially for TEM and FIBSEM, to about 10-20 microns [17, 61]. This cannot be judged easily by visual inspection of the sample alone. We found that XRM could be used for this purpose. Given that this is a non-destructive process, this allows for subsequent targeted EM image acquisition.

While several electron-dense probe types are available, we focused on miniSOG [50] and for proof-of-principle studies, labelled olfactory neurons in the *Drosophila* adult brain. We find that miniSOG labels can be targeted to distinct neuronal sub-compartments, such as membranes, cytosol, synapses and mitochondria. Through modifications in osmium and heavy metal staining, contrast introduced by the genetic label and on membrane boundaries, organelles and synaptic features can be further enhanced. The miniSOG label can be detected under TEM, SBEM or FIBSEM modalities. We explored the use of computer-based strategies to segment labelled structures throughout the volume and reveal some of the cellular and neuronal features that can be identified using these methods. Given the compatibility in contrast detection, we explored a new imaging technique, micro-computed tomography (micro-CT or X-Ray Microscopy/XRM). This allowed us to perform non-destructive X-ray imaging on intact brain samples to screen for optimally stained specimens. At submicron resolutions, macro- and micro-scale features, such as neuropil regions, axon tracts and individual cell somata could also be distinguished from these XRM scans. This was followed with targeted volume EM acquisition to investigate features of interest at nanometre-scale resolutions.

## Results

### Design, expression and label detection of miniSOG probes

Our first aim in probe design was to ensure miniSOG labels can be detected using fluorescence microscopy and that their expression did not cause morphological artefacts in the neurons they were expressed in. We cloned miniSOG DNA into *Drosophila* expression vectors and initially analysed probes in cultured *Drosophila* S2 cells under fluorescence microscopy (Figure 1A-C). We found that miniSOG fluorescence photobleached quickly under wide-field fluorescence illumination. To circumvent this, bright, photostable FPs (tagRFP or mKate2) fusions were incorporated. Using the *Drosophila* UAS-GAL4 system [62], their expression in the adult brain were monitored by confocal microscopy detecting either miniSOG or FP fluorescence or by immunostainings (Figure 1E-I). Several miniSOG probes were tested in this way, targeted to the plasma membrane, synapse, cytosol or mitochondria. Based on light-level analysis, we verified that their expression did not cause any gross axonal or dendritic targeting defects.

**Figure 1:**
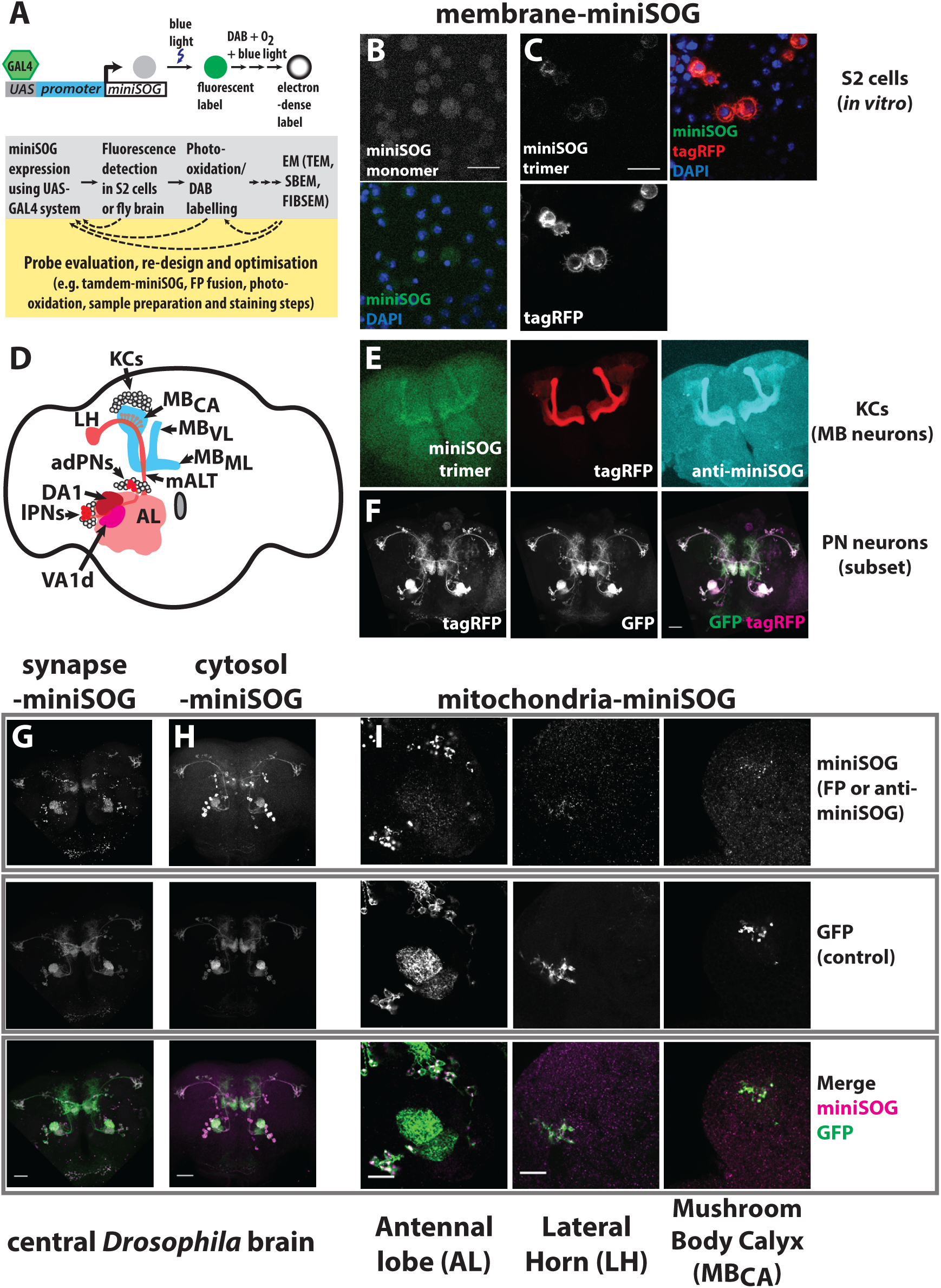
Design, expression and detection of miniSOG under fluorescence microscopy. - A-C. Summary of the design process adopting miniSOG as a genetically encoded electron-dense marker for *Drosophila* cell labelling (A). UAS-GAL4 expression system was used to express miniSOG gene product. Expression products were first tested in transfected *Drosophila* S2 cells grown *in vitro* (B, C). Membrane-targeted miniSOG (membrane-miniSOG) probes were expressed either as a monomer (B), or as trimer fused to tagRFP (C). S2 cells were imaged for miniSOG fluorescence, RFP or nuclear DAPI fluorescence, as indicated. Confocal images for miniSOG fluorescence was collected over high gain and extended pixel dwell time, resulting in higher backgrounds. Despite this it was not possible to gather enough signal to detect monomeric miniSOG as higher illumination led to photobleaching. Note the fluorescence in B is background and non-specific. Scale bars: 20 microns. - D. Schematic of mushroom body neurons (MB, also known as Kenyon cells or KCs) and a subset of projection neurons (PNs) in the *Drosophila* adult brain. These structures are bilaterally specified but only the left hemisphere is illustrated. From anterior dorsal (ad) or lateral (l) cell body locations at the Antennal Lobe (AL) periphery, most PNs have dendrites that densely innervate single glomerular locations. Projecting posteriorly along the medial AL Tract (mALT) to dorsolateral locations, PN axons have long-range projections that terminate at the mushroom body calyx (MB_CA_) and lateral horn (LH)[95, 96]. At the MB_CA_, PN axon terminal boutons connect with KC dendrites [72, 73]. From posterior dorsal locations, KCs extend axon anteriorly and bifurcate to form the vertical lobe (MBVL, also known as dorsal or alpha lobes) and medial lobe (MBML, also known as horizontal, gamma or beta lobes). The UAS-GAL4 system [62] was used to express miniSOG products in these neurons. The OK107-GAL4 labels KCs [97]. Mz19-GAL4 was used to label PN subsets, which innervated glomeruli subsets (DA1, VA1d, DC3) within the AL [90]. DC3 is not visible from the AL surface. - E. membrane-miniSOG expression in KCs, visualised under miniSOG or tagRFP fluorescence, or by Alexa® 633 secondary antibodies that cross-react to anti-miniSOG. Despite high non-specific background, antibody labelling correlates well with miniSOG and RFP. - F. membrane-miniSOG expression in a subset of PNs, using Mz-GAL4. As a control, a membrane-GFP reporter was co-expressed to determine the extent of overlapping projection patterns with the miniSOG probe. - G-I. Synapse (G), cytosol (H) and mitochondria (I) targeted miniSOG expression in PN cells were verified by fluorescence microscopy. miniSOG patterns were visualised either by immunostaining using miniSOG antibodies linked to Alexa**®** 568 (H,I) or through the fluorescent mKate2 fusion protein (G). This choice depended on the designed probe and expression levels detected. As a control for expression pattern, these brains also co-expressed membrane-bound GFP. Images were acquired from the brain region indicated next to each panel. Scale bars: 40 microns

As previously reported, miniSOG labelling is based on photo-oxidation [50] where blue light illumination stimulates the production of singlet oxygen that is used to oxidise the substrate, Diaminobenzidine (DAB). This results in a brownish, osmophillic stain, visible in brain regions where miniSOG was expressed (Supplementary Figure 1), serving as a useful indicator of the photo-conversion process. In our tests, specimens that expressed multi-copy versions of miniSOG, either as tandem motifs within a single fusion protein or multicopy transgenes showed stronger DAB labelling.

### Effect of contrasting agents on miniSOG-DAB labelling under EM

Having successfully introduced the probe into specific neurons by genetic targeting, we next determined whether miniSOG contrast could be detected under EM. Photo-oxidised samples were initially processed using standard osmium staining, followed by epoxy resin embedding. Ultrathin sections from brain samples that express membrane-targeted miniSOG (myr-miniSOG) in Mushroom body (MB) neurons (also known as Kenyon cells or KCs) or a subset of projection neurons (PNs) were imaged under TEM (Figure 2B-F). Label contrast was visible in superficial (alpha lobe tip, ~<10 micron from brain surface) and deeper layers (peduncle, ~100 microns from surface; mid-brain) of the brain (Figure 2B-C). Consistent with membrane localisation, the miniSOG-DAB label was localised to the membrane and proximal regions of the cytoplasmic space.

**Figure 2:**
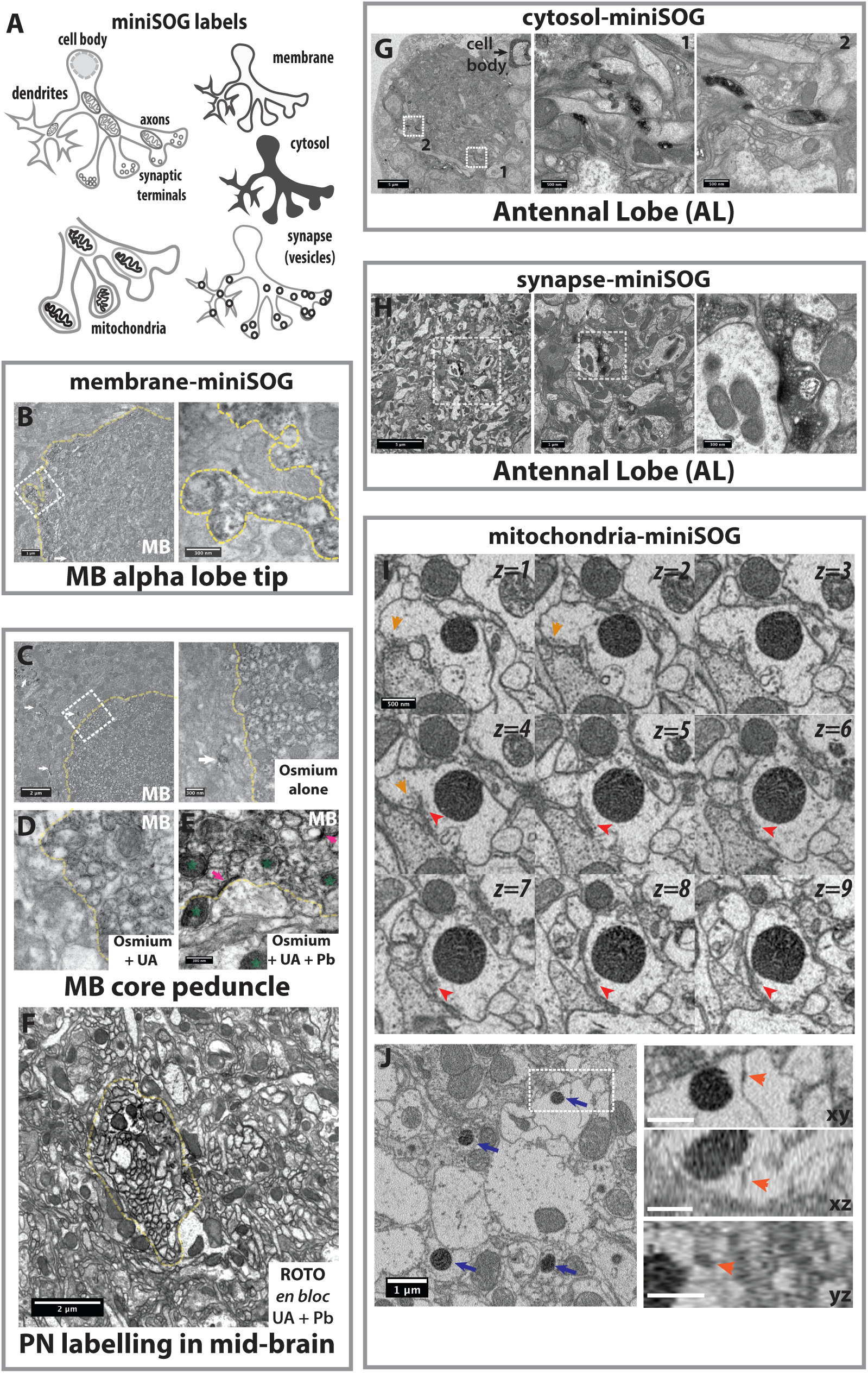
miniSOG targeted labelling and contrast detection under EM. - A. Schematic of the different miniSOG probes used to generate electron-dense signals in distinct neuronal sub-compartments. - B-C. TEM micrographs of osmicated, epoxy-embedded samples expressing membrane-miniSOG in MB neurons, sectioned at the vertical (alpha) lobe tip (B) or the MB peduncle (C). Higher magnification micrographs are shown on the right, from inset regions outlined (white dashed box). The miniSOG label is detected as electron-dense DAB precipitates on the membranes and inner cytoplasmic space. The dashed yellow lines indicate the border regions where the MB neuropil is present (MB). Next to the MB core peduncle, other neurons also express miniSOG (arrowhead). These non-Kenyon cells form part of the OK107-GAL4 expression pattern. Scale bars: 1 or 2 microns (B and C, respectively) or 300nm (insets). - D-E. The effect of applying contrast on-section using uranyl acetate (UA) and lead citrate (Pb) stains. Adjacent thin sections from the same osmicated, epoxy-embedded specimen as in C were used here. Note the enhanced contrast on the DAB and non-DAB labelled regions when UA with Pb stains are used together: on membrane borders, synaptic densities (magenta arrows) and organelles (green stars highlight mitochondria) and cytosol. Scale bars: 300nm (D, E) and 2 microns (E). - F. TEM micrographs showing membrane-miniSOG labelling of PNs (using *GH1469GAL4,* see Methods), sectioned at the mid-brain. Whole brain samples were osmicated following the ROTO protocol, together with *en bloc* UA and Pb stains. High contrast labelling of PN neuronal membranes is evident from the image. Yellow dashed lines indicate the PN neuropil boundary. Scale bar: 2 microns - G-H. Sectioned from the AL, TEM micrographs showing cytosol (G) or synapse (H) targeted miniSOG in labelled PN dendrites (using *Mz199GAL4*, see Methods). The specimen in G (Sample-*E1*) was treated with ROTO only. The labelled cell body on the left panel corresponds to a single adPN cell on the right AL. The sample in H (Ref: *S391*) was treated with ROTO, together with *en bloc* UA and Pb. Synapse targeting was achieved using a Synatotagmin-miniSOG fusion protein; hence the electron-dense contrast on presynaptic and vesicular structures. Scale bars: 5 micron, or 500nm (insets for G), or 1 micron or 300nm (H), as indicated in panel. - I-J. SBEM images of mitochondria-targeted miniSOG in PN axons (using *Mz199GAL4*), acquired from the LH (Ref: *1408079R04,* see Text). Two examples are shown. In I, consecutive *xy* sections of a single labelled mitochondrion in the centre are shown. Each *z*-numbered section is subsequently spaced 50nm apart. Synaptic features (vesicles and presynaptic densities; orange and red arrowheads, respectively) were found in close proximity to the mitochondrion. - In J, several mitochondria-miniSOG are visible (blue arrows). Highlighted by the box region and shown on the right are multi-planar (xy, xz and yz) views of a putative gap junction (orange arrowheads) that is next to a labelled mitochondrion. This specimen was treated with ROTO with *en bloc* UA and Pb. - TEM and SBEM images were obtained from transverse sections. For illustration purposes, image-processing filters was applied (Thresholding, Gaussian Blur and Sharpening, as necessary) in most images except for G and H where only Thresholding was applied.

Differences in osmium and heavy metal staining methods lead to variations in the labelling of cellular and tissue structures, affecting contrast and image detection on the electron microscope used [63–68]. Cytoplasm, membrane boundaries, organelles and synaptic densities can be enhanced by contrast stains. While on-section treatments (uranyl acetate and Reynold's lead citrate) for TEM is routine, this procedure is highly labile [64]. For volume imaging, this needs to be tightly controlled so that all sections achieve consistency. For fly brain samples, while on-section stains enhanced the miniSOG contrast, the use of *en bloc* (bulk) stains provided a good alternative as contrast on the miniSOG-DAB and the surrounding ultrastructure was also enhanced (Figure 2D-F).

Furthermore, *en bloc* procedures are not only applicable for block-face imaging (SBEM and FIBSEM), but they also help to reduce charging effects by grounding the sample with heavy metals [67]. Even with Reduced Osmium-Thicarbohydrazide-Osmium (ROTO) staining alone, good levels of contrast can be detected when imaged by TEM, SBEM and FIBSEM (Figure 2G; also Figures 3 and 4).

**Figure 3:**
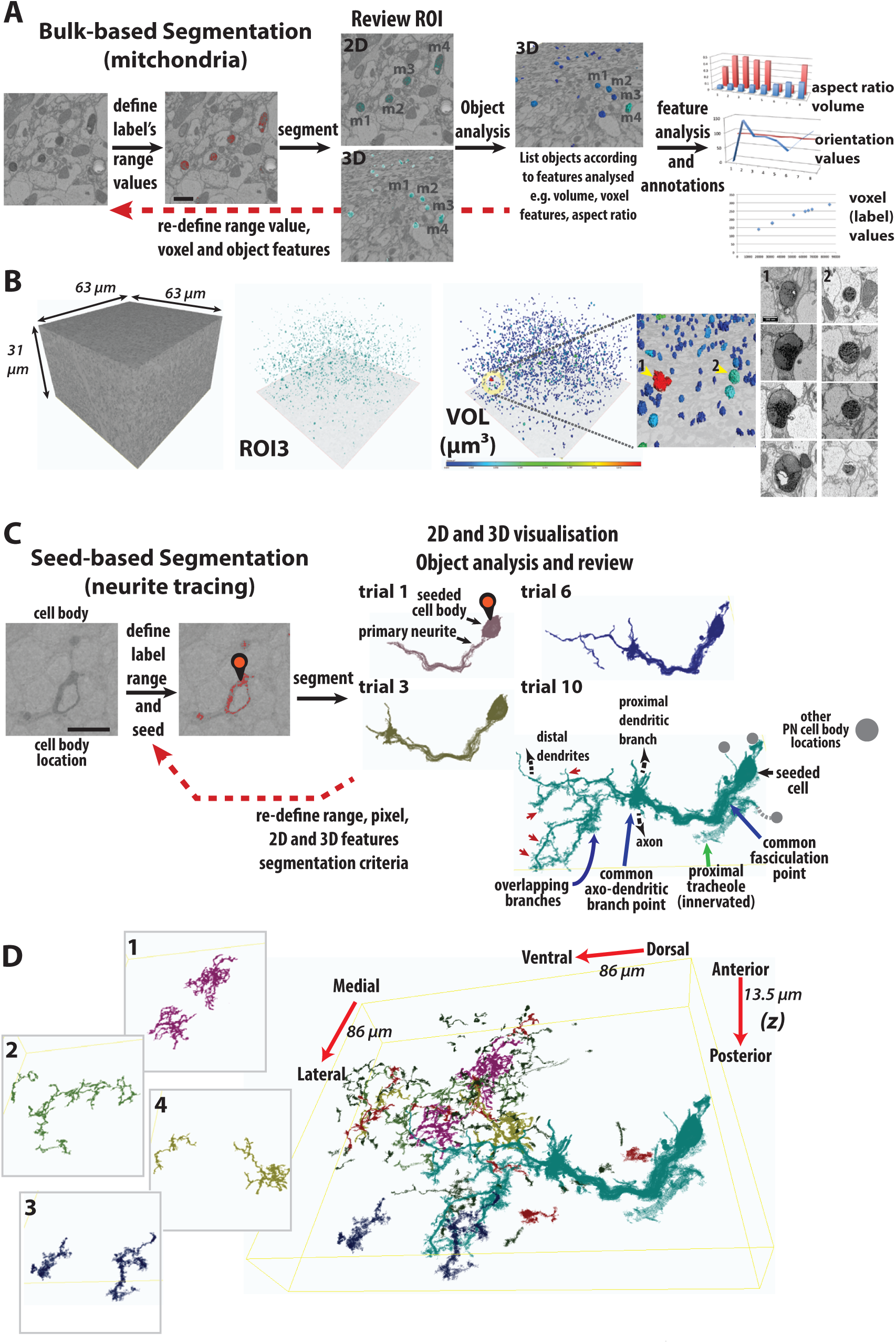
EM volume acquisition and computer-assisted segmentation of miniSOG. - A. Following SBEM volume acquisition, two computer-assisted segmentation approaches were tried. For the mitochondria-miniSOG labelled volume, given that the label was well contrasted and segregated, *bulk* segmentation was applied. Range values corresponding to the miniSOG contrast was determined from pixel probe readings of labelled mitochondria and interactively setting minima and maxima threshold values. As indicated by the red pixels, the selected range correlated with a subset of labeled mitochondria. However, all pixels corresponding to the desired range will be segmented from the image volume (ROI1, not shown). An Object Analysis step will list all the objects identified, together with their analysed features (voxel, volume and vectorial properties). Further filtering steps were used to restrict the desired ROI only to miniSOG-labelled mitochondria. High contrast but low volume objects were eliminated from the segmented dataset as these were corresponded to non-specific background signals (cutoff: <10 voxels; <0.0002 micron^3^). A further filter based on the variance intensity, eliminated larger, cell-based structures, such as highly contrasted cell membrane segments. The segmented objects presented from this set (ROI3, cyan-green) were then examined in 2D and 3D, comparing them against the underlying ultrastructure. Following this, an Object Analysis function was applied. This produced a mesh and listed all the segmented objects identified, together with their features (voxel intensities, volume and orientation properties). The volume mesh for a subset of mitochondria (m1-4) is presented in 3D view. Note the graphs on the right are for illustration purposes only. The SBEM image volume (Ref: *1408079R04*) was acquired from a specimen (*Sample 10*) that has been treated with ROTO with UA and Pb stains. Scale bar: 1 micron. - B. A summary of the mitochondria-miniSOG labeled objects found within the acquired volume (63×63×31 micron; voxel dimension: 18.9×18.9×50 nm). Post-filtering, the desired dataset (ROI3, cyan-green) can be visualized in bulk 3D. Object Analysis is used to set a criterion, here displaying objects according to size (VOL, μm^3^). The individual objects are size-indexed blue–red, from the smallest (0.0043 μm^3^) to largest structures (0.2785 μm^3^), respectively. Volume analysis allowed us to investigate the morphological basis of a single mitochondrion (1, red) that appears much larger from the rest within the dataset. Closer ultrastructure inspection (right panel) suggests it is undergoing breakdown. By comparison, an adjacent smaller mitochondria (2, cyan) retains has the expected pattern of labeling as shown previous *in vitro* studies [47]. See also Video 2. - C. In the second approach, we used a *seed9*based segmentation method to trace labeled neurites. This was applied to cytosol-miniSOG labelled PN neurons as that the label appeared contiguous and thus likely to have overlapping voxels (see Video 3). After defining the range values, a seed point (black pin icon) was placed at the cell body location. A Point and Click seeding function initiates the 3D segmentation process. Several trials were performed based on a threshold range of pixel intensities. Set to common maximum, increasingly lower range intensities were sampled in subsequent trials (trial/ROI 1-12). Each step change represent 0.2-0.4% of the true range. Segmentation at trial 1 (ROI1) traced primary neurites that form part of the seeded cell body. Subsequent trials show the segmented results (ROIs) with increasingly ramified structures. Some of these segments trace back to the cell body seeding point. They also included very thin protrusions, which emanate from dendrite segments or primary branches (red arrowheads). However, other traces represented overlapping segments from other labeled PNs. These ‘crossover’ traces occur at distinct locations where several labelled PNs have common cell body locations, branching points or overlapping branch segments (blue arrows). Within this volume, we found the target PN innervated a trachiole (green arrow). Given its proximity and elevated contrast within this structure, this too becomes incorporated to ROI10 when low threshold values (trial10) were used. - Two distinct events take place as increasingly lower values are sampled. Distal branches and terminal structures are included to primary branches. However, non-related, overlapping neurites, cell bodies and highly contrasted structures can also become incorporated into the primary cell process. See also Video 3. - SBEM image volume (Ref: *1411279R01*) was acquired from a specimen (Sample-*E1*, as used in TEM imaging and shown in Figure 2G) that was treated with ROTO with UA and Pb. The volume image corresponds to mid - posterior location on the right AL. For segmentation analysis, the entire stack (85×85×13.5micron) was image pre-processed by applying non-local means (sigma=5). Volume stack was downsampled (3×3×1) with a voxel resolution of 31.5×31.5×25nm. - D. Following the initial cell body tracing, additional traces were performed with the AL sub-volume, using the same seeding strategy and threshold values based on trial10. By reviewing on a section-section, multi-planar basis, seed points were placed on potential neurites. Many additional neural processes were found. A subset of these larger examples (1-4) is highlighted on the left. They have been highlighted in different colours to ease their visualization. A composite of these, together with smaller segments (dark green) and the PN cell (ROI10) are shown on the right. Many of these segments are predominantly located at the edge of volume or at distal branches regions (particularly for smaller segments). See Video 4 and Video 8.

**Figure 4:**
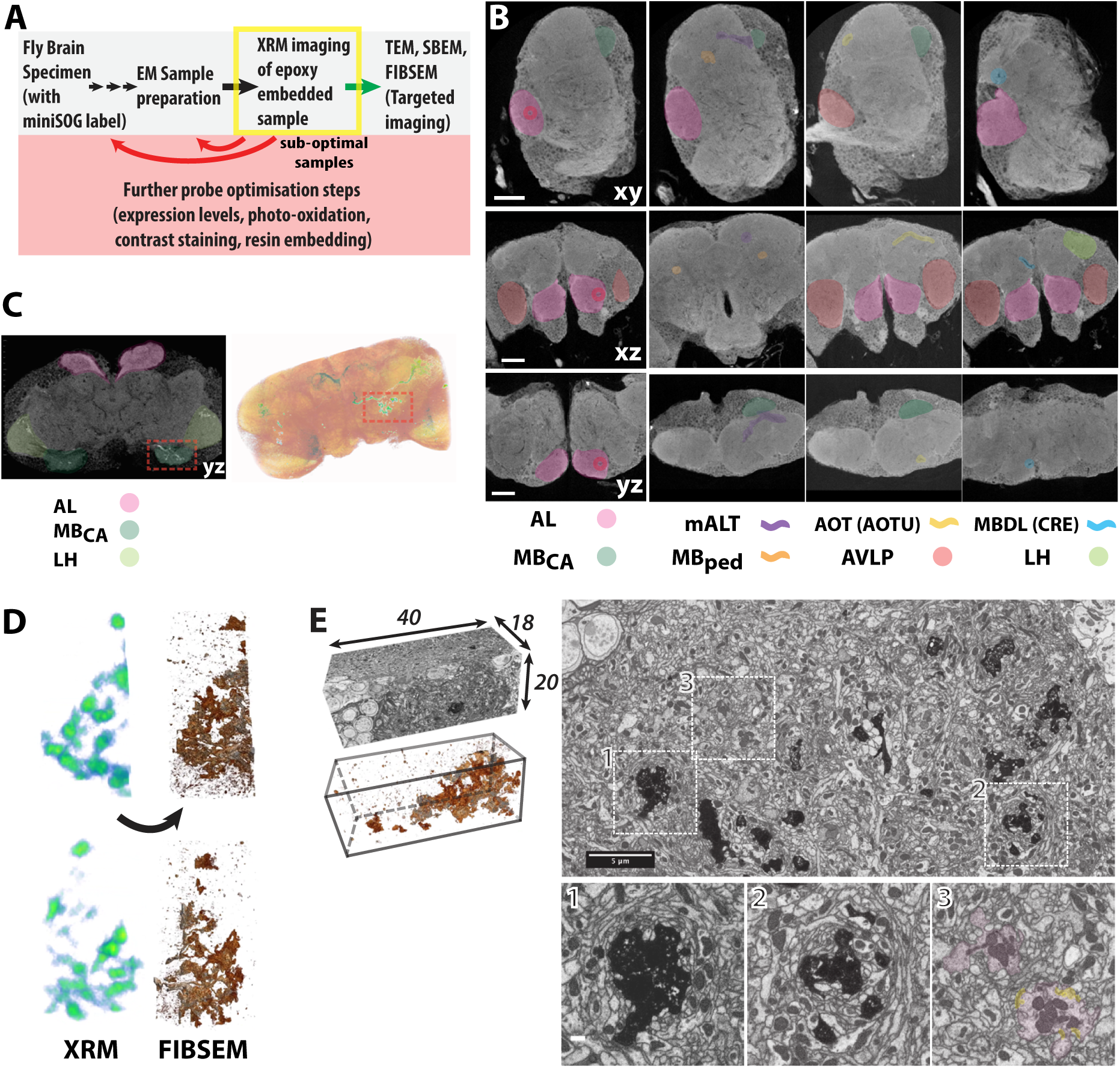
Whole fly brain miniSOG detection with correlative XRM-FIBSEM imaging. - A. Schematic of the XRM-EM workflow used to screen fly brain samples optimized for heavy metal contrast and for targeted volume acquisition. - B. XRM images were reviewed on a 2D section-section basis from all plane (xy, xz and yz) perspectives. Many notable neuroanatomical features were discernable. Neuropils highlighted include the AL, MB_CA_, and anterior ventrolateral protocerebrum (AVLP). Fasicles include MBped, mALT and axons within the AL (red circle). The Anterior Optic tract (AOT) and the Median Bundle (MDBL) are also visible and within in the regions of the Anterior Optic Tubercle (AOTU) and Crepine (CRE), respectively. Some of these were previously seen under light microscopy [70]. - C. Left, A single virtual XRM section image of a fly brain specimen with cytosol-miniSOG labeled PNs. Shown from the yz plane, the AL, MB_CA_ and LH locations are highlighted. Highlighted regions include AL, LH, and MB_CA_. Right, 3D volume rendered image of the entire specimen. When thresholded, the miniSOG contrast in PN axons is immediately recognisable, appearing in green and red, based on a blue-red color ramp for low-high signals. Note the uneven PN labeling on the left side of the brain. As such, the right MB_CA_, boxed in red, seems the most appropriate region for FIBSEM volume acquisition. - D. Left, Correlated miniSOG images from XRM (green-red) and FIBSEM (as copper-silver for low-high signal intensities) tomograms, shown from two perspectives. - E. Left: acquired FIBSEM volume of MB_CA_ (dimensions is indicated in microns; approximate voxel size is 10nm isotropic). As shown below thresholding reveals the miniSOG label. Right: Single xy plane section. The panels below are higher magnification from the inset regions, as numbered. The very high contrast, appearing as black, corresponds to miniSOG labeled PN bouton terminals. A single non-miniSOG labelled PN terminal is highlighted, in pink, for comparison. Due to technical difficulties, it was not possible to obtain higher resolution of synapses in this specimen. However, synaptic components, such as densities and some vesicles are discrenable, highlighted in yellow. This specimen was treated using the ROTO only protocol.

### miniSOG contrast detection in other neuronal sub-compartments

We generated miniSOG probes targeted to cytosol, synapses or mitochondria that produced a good level of contrast in these subcellular compartments (Figure 2G-J). These probes are useful in several ways. First, synapses and mitochondria are highly relevant to neuronal function and are best studied at nm-resolution: genetic labels can be used to visualise them with cell-targeted precision. Second, large numbers of labelled structures can be selectively visualised in very large volumes (see Figure 3). Third, some probes can act as proximity markers, by labelling relevant neurons or associated structures without obstructing other relevant ultrastructures. As illustrated in Figure 2I and J, putative synapses and gap junctions can be distinguished close to mitochondria-labelled miniSOG. Last, acting as a whole cell tracer, the cytosol-targeted miniSOG has been useful in outlining labelled neurons (see Figure 3 below).

### Enhanced contrast based segmentation from miniSOG-labelled EM volumes

We next generated EM volumes from miniSOG-labelled samples. Different acquisition and segmentation strategies were explored to see how to best visualise the label from image stacks. From the *Drosophila* brain, images were collected from the different regions of olfactory processing centres [69], such as the lateral horn (LH, or lateral protocerebrum) that is a higher order olfactory processing centre and target site of PN axons (Ref volume: *1408079R04*; dimension (x/y/z): 63×63×31 microns; voxel dimension: 6.3×6.3×50nm; number of z-sections: 624) and the Antennal Lobe (AL), which is the olfactory processing site between olfactory sensory axons and projection neuron dendrites (Ref volume: *1411279R01*; volume dimension: 86×86×13.5 microns; voxel dimension: 10.5×10.5×25nm; z-sections: 541). These image stacks were acquired on two SBEM microscopes that have different sensitivities and configurations (see Methods). Yet, robust contrast on miniSOG labels and the surrounding ultrastructure (membranes, organelles and synaptic compartments) was visible from both image volumes (Figure 3, see Videos 2 and 3).

Given the enhanced contrast, segmenting miniSOG-DAB labelled structures should be straightforward. While manual segmentation is the most direct, we focussed on automating segmentation for several reasons. First, computational tools can readily identify labelled structures from the image stack by thresholding. This was the case when *bulk* segmentation was applied to a SBEM volume that had mitochondrial-targeted miniSOG (Figure 3A and B, Video 2). Second, once defined, such regions of interest can be further interrogated computationally for desired features of interest. For instance, we can determine pixel values, volume, aspect ratio and orientation properties of the segmented objects (Figure 3A, Video 2). Third, we find computer-assisted visualisation is able to detect subtle differences in the contrast due to miniSOG labelling. This was particularly useful when applied to low contrast-low signal images, as shown in the seed segmentation example, thus achieving greater accuracy in neural tracing (Figure 3C and D, Videos 3 and 4). Fourth, and related to the above, computational tools will have a robust record of the selection criteria used. This is useful if the procedure needs to be replicated across different datasets, or altered to define, correct or specify for new or existing features or sub-features. Last, computational tools can readily exploit 3D segmentation tools in the first instance. This aids quicker review over the 3D objects identified (Figure 3).

Note however that while initial results from such segmentation strategies can be obtained relatively quickly, cell biological and neuroanatomical expertise is necessary to review the identified objects. Following initial review, iterative segmentation steps help to refine the desired objects (Figure 3, see Discussion). As these probes can also be visualised under fluorescence microscopy, EM traces can be cross-validated with the light-based images.

### Micro-CT/XRM visualisation and correlation to brain regions for targeted EM volume acquisition

We were motivated to test micro-CT/XRM as an application to, 1) select optimally stained samples for EM data acquisition (prior to lengthy EM volume imaging), 2) for its ability to image whole brains non-destructively and, 3) to obtain cell biological and neuroanatomical information across very large volumes at sub-micron resolutions (Figure 4A). Osmium-treated and epoxy-embedded fly brains were imaged on a Zeiss Xradia Versa 520 lab-based scanner. From these scans, many features were recognisable. Images were similar to those obtained from light microscopy and from scans obtained using synchrotron-based X-ray sources [[70]][71], respectively; Figure 4B]. Our XRM images also highlighted individual cell somata from the cortical rind (perikaya) surrounding the brain, as well as axon tracts. Both appear negatively stained. Surrounding cell membranes and densely packed neuropil regions such as the antennal lobe (AL), mushroom body peduncle (MB_ped_) and calyx (MB_CA_) and optic lobe lamina and medulla layers showed higher levels of staining (appearing white or light grey in Figure 4B, see also Video 5).

A recent study showed miniSOG generated contrast in mouse brain tissue can be detected using XRM [59]. When miniSOG labelled *Drosophila* brains were scanned, contrast from labelled PN neurons clearly highlighted axon tracts and terminal boutons (appearing white in Figure 4C; Video 6). However, labelling from within the AL glomeruli was not seen. One possible reason for this is the high level of native contrast in this densely innervated region makes any genetically introduced contrast less likely to stand out.

As part of the EM workflow, we transferred the miniSOG labelled sample to a FIBSEM microscope. From the initial XRM scan, we noticed that the bilateral miniSOG^+^ PN axonal structures were unevenly labelled within the specimen, with the left side having lower signal at the PN terminals in the MB_CA_ (Figure 4C). The reason for this is unclear, it may have occurred during the photo-oxidation or the post-staining process. Nonetheless, the XRM scan allows us to select the MB_CA_ on the right hemisphere as the ideal region of interest from which to acquire a FIB image sub-volume, which was approximately 0.075% of the total brain volume.

We next established a correlation between FIBSEM and XRM by registering the SEM field image on the sample surface to virtual slices from the XRM volume data, enabling precise targeting of the region of interest. Using FIBSEM, we acquired image stacks from the mushroom body calyx (Ref volume: *1509159 R2/5*; volume dimensions: 40×20×18 microns; voxel dimension: approx. 10×10×10nm; z-slices: 1800). Here we find the overall contrast and of the level of miniSOG labelling of the cytosol of PN axon terminals were very high. Comparisons between the XRM and SEM tomograms showed the label contrast closely matched each other (Figure 4D; Video 7). We are interested in the Mushroom Body calyx (MB_CA_) as it is a higher order olfactory processing region where PN axon terminals synapse with Kenyon cell (KC) dendrites [69]. Indeed, closer inspection of the FIBSEM images revealed very fine diameter KC dendrites surrounding the PN boutons (Figure 4E), similar to previous reports [72, 73].

In summary, using XRM, we were able to screen candidate samples for optimal labelling characteristics and perform site-specific volume image acquisitions using FIBSEM. Having whole brain XRM images also provides many cell and neuroanatomical relevant features that is useful for subsequent reconstruction and analysis.

## Discussion

Probe labelling is a key method in biological imaging. The ability to highlight specified structures helps decipher how macromolecular complexes, cellular components, cells and tissues are organized, giving insight into their function. In neuroscience, given the diversity of molecular, cellular and physiological substrates, fluorescent probes have been used extensively in cell biological, neuroanatomical and neurophysiological studies [74–78]. Yet adopting them for the purpose of light-based neural circuit reconstruction still has limitations as it is unclear how segmentation can be implemented on densely labelled neurites across large dimensions and at high resolution. Volume EM techniques are therefore favoured, with acquisition volumes of >6,000,000 µm^3^ and 4-50nm voxel resolution already reported [19]. However, given the large amount of high-density data, working out how individual segments of axon, dendrites and synapses are represented as neural network-based entities requires substantial reconstruction time [15].

Hence, there is a huge attraction in applying labelling (particularly correlative) methods to volume EM datasets to enable targeted reconstruction. Indeed, many innovative approaches are available. Apart from those already mentioned above, other probe types include antibodies, FPs, fluorescent dyes and nanoparticles [5, 79–83]. While each technique is different, they all introduce variability associated with sample handling, labelling, image alignments, and resolution mismatches. Trade-offs are inevitable when balancing the requirements for ultrastructural preservation and EM contrast with correlative fluorescence visualisation [84]. These factors make it challenging to consistently generate high contrast EM image volumes with labelled features of interest.

In this study, we have focused on EM-visible probes, which are genetically-encoded, highly compatible with sample preparation methods and capable of high resolution visualisation. These criteria facilitate ease of labelling, promote ultrastructure preservation and contrast generation. This allows a single EM modality for label detection and cellular ultrastructure discrimination and both are acquired at identical nm-scale resolutions. Our feasibility study acquired volume datasets from olfactory centres in the *Drosophila* adult brain that are densely innervated. From these studies, several observations were made, which are discussed below.

### miniSOG probe expression in neuronal cells

When expressed in *Drosophila* neurons, we find that miniSOG works effectively when targeted to distinct cellular sub-compartments. This is a marked improvement over other EM protein-based probes such as HRP, which have limited targeting capabilities [46, 48, 53, 55], or ReASH, which necessitates cell-toxic conditions for probe introduction [85]. One initial concern was whether the inaccessibility of blue light in deeper tissue regions might hamper the labelling process [47]. With protocols used here, DAB label was visible in the mid-brain, at ~100 microns depth. One protocol adaptation was to perform photo-oxidation using blue-light LEDs (see Methods). One area of continued development will be to gauge how much DAB label is sufficient for image detection. For example, our current approach relies on a tandemisation approach: attaching multiple miniSOG probes to a single targeting motif. The samples used also carried 1-4 transgenic copies for expression. While this does not impact on the study's conclusions, ideal probes ought to be small and minimally expressed in order to reduce the possibility of disrupting cell function and morphology. In this respect, improvements may come from re-engineered probes or using related probes to increase DAB labelling [86, 87]. Related to this, our fluorescence imaging tests suggest that not all probes tested are currently optimal. For example, while the mitochondrial-targeted miniSOG does not perturb PN axon or dendrite targeting, as judged by light microscopy, mitochondria labelled miniSOG appeared aggregated at the cell body and less frequently in distal axon termini. Closer EM-level examination shows some mitochondria appear smaller and fragmented. At this resolution, axon fragmentation was also prevalent in the LH in the specimen of a 7-day old adult fly. This indicates high levels of mitochondria miniSOG expression may lead to neurotoxicity over time. Related to this, expressing high-levels of miniSOG in the mitochondria of *C.elegans* reportedly caused neurotoxicity and cell death [88]. Nonetheless, given the precedent that label contrast can be detected at mitochondrial locations and serve an important purpose, refined probes will facilitate further improved use.

### miniSOG and ultrastructure contrast detection under various EM modalities

Labelling probes should fulfil several key criteria. Exhibiting highly localised contrast is critical. The label needs to be detected uniformly throughout the regions and structure of interest targeted. Label generation needs to be minimally disruptive for image detection; any background artefact introduced has to be minimal. Several aspects were addressed in this study. Highly localized miniSOG contrast was consistently observed. Segmentation throughout the volume suggests artefacts and background effect were minimally disruptive. In any imaging paradigm, there are always concerns regarding the extent of targeting and labelling density any probe can achieve. In the course of neurite tracing using cytosolic miniSOG, we found that segmentation errors can result from signal drop-off, particularly at dendritic branch points and at large axonal segments where it was more difficult to achieve high or uniform levels of labelling. Such errors are easily detected when imaging at nanometre-scale resolutions. Nonetheless, with expert knowledge of projection patterns and by looking at the underlying ultrastructure, such errors can be easily rectified (see Video 8 for examples). Thus, one area of continued development will be to determine how each designed probe could achieve optimal labelling at targeted regions of interest. One easy way is to allow for more DAB label to develop. However, overdevelopment can result in the loss of resolution and generate labelling artefacts (Supplementary Figure 2). Masking, where the underlying ultrastructure is obscured by DAB labelling, is also another concern.

The choice of electron microscope determines the type of images that can be acquired [17]. It is encouraging that our results show all tested EM modalities yielded high-contrast miniSOG labelling. While the present study was not focused on maximising the resolution possible for each microscope, future work will optimise image acquisition parameters not only to detect the genetically introduced labels but also to ensure the desired ultrastructural components (particularly synaptic substructures) are well represented throughout the image volume. Related to this is the issue of ultrastructural preservation. While aldehyde fixation is the most common method, high-pressure freezing and freeze substitution protocols provides more native organisation of cell architecture, synaptic components and the extracellular space [89]. Future work will determine how this can be incorporated within the labelling workflow.

### Label detection and segmentation strategies using computational approaches

Labelling approaches provide a simple way to extract information from complex biological images. Given the elevated contrast introduced by miniSOG, labelled neuronal components can be easily identified from high-density EM images. Computational tools work well in this respect as discrete pixel-based segmentation criteria can be applied to the entire volume. As low- and high-order features become attributed to each segmented object, this forms a valuable way of linking them within the volume, facilitating large-scale, comparative and interpretive analysis. Future work might explore how such tools work best and the types of information that can be obtained with each probe. More tools will inevitably be needed to complement and complete the reconstruction process. This might include semi-automated and rules-based segmentation algorithms that reflect neuronal morphologies and cell-based structures. While this manuscript was being finalised, we were encouraged to find similar approaches were also being developed to exploit genetically encoded EM marker based neural tracing in mouse neurons [49]. Similar approaches to adopt unsupervised segmentation algorithms of labelled neurons will be useful in our future work.

In our seed segmentation trials, it is clear that identifying the PN cell body and using its location as the seed point is insufficient to generate an accurate trace pattern throughout the acquired volume. In contrast to well-segregated, mitochondria-miniSOG labels, neurite tracing using cytosolic-miniSOG showed a high degree of under-segmentation. One limitation in our samples was that several overlapping PN neurons were genetically labelled [90]. Thus, depending on the probe, its labelling pattern and the contrast features within the volume, more sophisticated segmentation tools will be needed. Further genetic tools can also be used to prevent overlaps, by restricting targeted expression patterns.

One interesting property of computer-assisted segmentation is the high degree of morphological complexity found in the dendritic arbours of labelled PNs, particularly in the lateral regions (Example 3 in Figure 4D). The organisational significance of these intricate branch protrusions is unclear. Interestingly, high-resolution analysis in the AL of silkmoth *Bombyx mori* also show similar dense protrusions from the dendritic shafts of PNs [91]. These features are also reminiscent of the ‘twig-like’ or ‘spine-like’ structures recently found throughout *Drosophila* neurons analysed, where thin actin-rich and microtubule-free terminal neurites show high levels of synaptic input [92]. Future work will determine whether these tufted protrusions on PN dendrites carry specific molecular and regional identities or synaptic patterns required for AL connectivity.

### Correlative XRM-EM volume imaging of the fly brain

Many aspects of fixation, labelling and contrast generation are not well understood and have led researchers to reinvestigate EM sample preparation methods to achieve better consistency [63, 66, 67, 89, 93, 94]. The inability to predict if EM specimens are optimal leads to wasted efforts, which on any given microscope incurs days-weeks of acquisition time. If sub-optimal image volumes become incorporated into the segmentation workflow, this adds to further waste. This prompted us to investigate XRM as a method for checking sample quality. Our initial testing has shown it to be critical prior to EM volume acquisition. In addition, XRM has become an invaluable imaging tool, revealing cell body and long-range projections patterns as well as outlining discrete brain regions. Future work will investigate how the XRM imaging can be used as a part of a reconstruction workflow to trace neurons across scales of whole brains, brain regions, axon and dendrite projections and ultimately leading to synaptic connectivity patterns under EM.

### Future perspectives

In summary, our study explores the implementation of EM-based reporters to label neurons in a whole brain setting. We propose a set of tools that use genetically-encoded probes, enhanced contrast staining, automated image volume acquisition and computer-assisted segmentation for the dissection of complex neural circuits. By combining genetic labelling with the structural analysis of molecularly defined components, such targeted approaches help uncover local network motifs and synaptic connectivity patterns. They are especially valuable in enabling genetic targeting of identified neurons even within a restricted EM volume that does not include enough of the neuronal arbour for cell type identification by morphology. With enough examples from different samples and brain regions, their insights can contribute to wider EM neural reconstruction efforts that are taking place across larger regions and to the general understanding of the nervous system organization.

Although our own research focuses on neural circuit analysis, these tools will also be very useful in cell and developmental biology and disease model studies. With a focus on large dimension, high-resolution based imaging, EM probes can be used to identify rare features (e.g. when searching for transient structures or cell types that arise from morphogenetic or pathological events [60]), or to acquire large datasets for ultrastructural studies of specific cellular components.

## Methods

### *Drosophila* stocks and immunohistochemistry

Brain samples used were 3-7 day old adult males. GAL4 drivers used have been previously described [90, 97, 98]. Immunohistochemistry was performed as previously described [99]. miniSOG antibodies were generated in guinea pigs (Eurogentec S.A., Seraing, Belgium).

### miniSOG design, vector construction and Drosophila transgenesis

miniSOG DNA was custom synthesised (GeneArt, Life Technologies) for optimal codon expression in *Drosophila*, with the addition of restriction sites at the 5’ and 3’ ends to facilitate in-frame fusion cloning. To make tandem versions, several fragments were mixed together during restriction cloning into *Drosophila* expression vectors [100]. Probes were targeted to the plasma membrane using a Src myristoylation tag [100]. miniSOG tagging the *Drosophila* Eb1 protein resulted in cytoplasm localisation. *Eb1* DNA (*Eb19PE*; Flybase ID: FBpp0089088) was made synthetically, as above. Synaptic targeting was achieved by fusing miniSOG to Synaptotagmin, similar to previous strategies [99]. Mitochondria-localisation was based on miniSOG fusion to a *Drosophila* gene product CG18769 (Isoform PG; Flybase ID: FBpp0300165), based on its sequence homology to the Mitochondria Calcium Uniporter gene (MCU). Its labelling characteristics in *Drosophila* neurons under EM (Figures 2I and 3B) are very similar to previous results [47]. All miniSOG fusions were incorporated at the COOH-terminal to the above-targeted proteins. Standard molecular biology techniques were used to generate these plasmid products. *Drosophila* transgenic lines were derived from germline transformations (Bestgene Inc., CA, USA) using the ΦC31/attP system [101].

### miniSOG expression in S2 cells and *Drosophila* brain

miniSOG was initially tested in *Drosophila* S2 cells as described previously [102] and analysed under confocal microscopy. miniSOG probes were expressed *in vivo* using neuronal GAL4 driver lines to express UAS-miniSOG products in mushroom body Kenyon cells (OK107-GAL4 [97]) or projection neurons (PNs) [using the GH146-GAL4 [[95, 98] or Mz19-GAL4, that label PN subsets innervating glomerulus DA1, VA1d and DC3)] [90]. Fluorescence images for miniSOG, tagRFP and mKate2 fluorescence data were acquired as previously described [50, 103, 104].

### Sample preparation for miniSOG photo-oxidation

Brain samples were processed similarly to previous studies [50], with minor modifications. *Drosophila* adult whole brains were fixed overnight in 4% paraformaldehyde, 0.5% glutaraldehyde in 2mM CaCl_2_, 0.1 M Cacodylate, (pH 7.4). These samples were kept cold whenever possible. Fixed brains were rinsed in chilled cacodylate buffer and were incubated in blocking buffers (in 0.1M cacodylate) sequentially for 10-15 minutes each time in (1) 10mM Glycine, (2) 10mM Potassium Cyanide, (3) 5mM 3-aminotrizole and (4) 5mM Mersalyl acid, to reduce non-specific DAB staining. miniSOG photo-oxidations were initially performed using a 150W Xenon light source (Lambda DG-4alpha, Sutter Instruments, CA, USA) with a 20×/NA 0.75 Plan Fluor Nikon objective that was mounted on a Nikon TE2000 microscope. At 470nm, this was approximately 1W/cm^2^ with 3.8mm^2^ field of view. Subsequently, a LED-based photo-oxidation unit was built (CREE; Code: 2216679; Part: XBDROY-00-0000-000000L01). This achieved illumination stability where several samples could be processed simultaneously. Samples were photo-oxidised for 2-20 minutes, being kept at 4°C throughout. For some probes, it was possible to monitor the progress of this process due to the presence of the brownish DAB precipitation. This was not the case for mitochondria-localised miniSOGs.

### Brain sample preparations for osmification, resin embedding and image acquisition

Different osmium and heavy metal stains were applied, following previous protocols [63, 64]. The variations in osmification included the use of 1% osmium tetraoxide (standard), 1.5% potassium ferricyanide with 1% osmium tetraoxide (reduced osmium or RO) or reduced osmium-thiocarbohydrazide-osmium (ROTO; RO as above, followed by 1% thiocarbohydrazide and a second round of 1% standard osmium). For contrast stains, *en bloc* uranyl acetate and lead aspartate were applied to whole-mount samples. On-grid staining included uranyl acetate and Reynold's lead citrate; applied for 2-5 minutes prior to TEM imaging [64]. Epoxy (Durcupan ACM, Sigma Aldrich #44611) embedded samples were sectioned and visualised by TEM (Technai T12 Spirit, FEI, Hillsborough, OR, USA), using a 2K×2K camera detector (BM-Orius, Gatan, Pleasanton, CA, USA). Following TEM verification, miniSOG-positive specimens were imaged under SBEM or FIBSEM. SBEM data was acquired using a 3View serial-block-face imaging unit (Gatan, Pleasanton, CA, USA) installed either with a high-vacuum Merlin or Sigma variable pressure (VP) field emission-SEM (Carl Zeiss Microscopy, Jena, Germany), equipped with either a 32K × 32K or 8K × 8K camera detector, respectively. We noticed that SBEM operating at high vacuum will produce better contrast and resolved images than those operating under variable pressure mode. This is a direct result of absence of gas scattering and higher surface sensitivity enabled by lower beam energies required to acquire data during probe scanning. Similar results were recently reported elsewhere [33]. Apart from SBEM, volume data was also collected on an Auriga Crossbeam FIBSEM, equipped with the ATLAS 5 scan generator and software tools (Carl Zeiss Microscopy, Jena, Germany).

### Image processing, visualization and segmentation analysis

Confocal microscopy data were acquired and processed as previously [99]. For EM Images, individual TEM micrographs, single or sub-stacks of SBEM and FIBSEM images were inspected using Fiji [105]. Image pre-processing was applied as necessary to remove background noise, improve visualization and figure presentation. These filters included Thresholding, Smoothing, Gaussian blur, Non-Local Means and Sharpening. Image stack alignments were also performed as necessary using StackReg on Fiji. Figures are presented in 8-bit to reduce file size.

Large EM volume stacks were visualized and analysed using ORS Visual SI Advanced software (Object Research Systems Inc., Montreal, Canada; Carl Zeiss Microscopy GmbH, Jena, Germany). To facilitate better performance on the workstation, datasets were downsampled by binning 3×3×1 (resulting with voxel resolutions of approx. 18.9×18.9×50nm (*1408079R04*) or 31.5×31.5×25nm (*1411279R01*). Video editing was performed using ORS Visual SI Advanced and on iMovie (Apple Inc., Cupertino, CA, USA).

### XRM/micro-CT analysis of epoxy embedded Drosophila brains

X-ray microscopy (XRM) was performed using a Zeiss Xradia Versa 520 (Carl Zeiss X-ray Microscopy, Pleasanton, CA, USA). The following settings were used to collect tomographic images. The emission source was set at 70-80kV and 6-7W. The samples were placed 8.9mm from RA-Source and 6.4mm for RA-Detector. A LE1 source filter was used. A 40× optical magnification objective was used, resulting in a field image of 0.409mm × 0.409mm. With a 1950×1950 pixel image size, this translates to a 200nm voxel size. The estimated reconstructed spatial resolution is 700nm. The total scan time lasted 22-24 hours, based on 17 sec exposure time and 360° degree rotation and step size of 0.11 for a total of 3201 projections.

## Acknowledgements

We thank the following for their contributions: Jeremy Skepper and Lyn Carter (CAIC, University of Cambridge, UK) for initial help with EM sample preparations and TEM sectioning. Tom Deerinck and Mark Ellisman (NCMIR, University of California, San Diego) for help establishing miniSOG protocols. Lucy Collinson and Chris Peddie (Francis Crick Institute, London) for help with SBEM acquisition on the 3View-SigmaVP and advice on image processing. This was acquired under the auspices of a MRC research grant awarded to the University of York (UK) and Cancer Research UK London Research Institute; The EM Core Technology Facility at Lincolns Inn Fields, London, provided services and the authors acknowledge its funding from Cancer Research UK, and from the MRC, BBSRC and EPSRC under grant award MR/K01580X/1’. Chris Guérin and Anneke Kremer (VIB Bio-Imaging Core, Ghent) helped with SBEM data acquisition on the 3View-Merlin. Matt Wayland (University of Cambridge) and Joe Ceremuga (Zeiss XRM, CA) for help implementing ORS image processing. Jimena Berni, Lucy Collinson, Philip Schlegel, and members of the Jefferis and Landgraf labs provided comments on the manuscript. This work was supported by European Research Council Starting Investigator and Consolidator grants, the European Molecular Biology Organization (EMBO) Young Investigator Programme, and Medical Research Council (MRC) Grant MC-U105188491 (to G.S.X.E.J).

---

**Supplementary Figure 1: Effect of increasing miniSOG motifs on DAB labelling**

- A. Transmitted light images of *Drosophila* brains post-DAB labelling/photo-oxidation. This process can be monitored by the presence of DAB precipitate, which is seen in MB neurons, where membrane-miniSOG is expressed. The DAB labelling extended over a 5-20 minute period. This process is blue light dependent as samples kept in the dark or exposed to green light do not show these stains (not shown). Blue light exposure of control fly brain samples (*w^1118^*) also did not result in such patterns (not shown).
- B. Expressing multiple transgenes further enhanced DAB labelling.

**Supplementary Figure 2: Effect of DAB over-development *in situ***

- TEM micrographs, sectioned at the AL, showing synaptic vesicle labelled miniSOG in PN dendrites (expressed using *Mz199GAL4*). From a larger panel, three higher magnification regions are shown on the right. Note the effect of excessive photo-oxidation/DAB development not only results in higher contrast but also in localised artefacts in neighbouring structures such as in mitochondria (highlighted in pink), together with the appearance of tiny holes (blue) and tears (yellow arrows). These effects on mitochondria structure are probably due to free radical damage generated from photo-oxidation. Tears and holes may be from reduced resin infiltration and subsequent fragmentation during sectioning, hence the high electro-lucent contrast within. This specimen (Ref: *S194*) was treated with the ROTO only. The image corresponded to the anterior of the AL. Scale bars: 5 micron or 500nm (insets), as indicated in panel.

**Video 1: Volume rendered images of mitochondrial-targeted miniSOG from confocal microscopy**

Two video sequences showing fluorescent image stacks acquired from PN neurons that express mitochondria-miniSOG (magenta, by antibody staining) together with control myr-GFP (green), acquired at AL or MB_CA_ and LH locations, respectively, as shown in Figure 1I. Following a 2D section-section review, 3D volume renders were generated from these stacks. Thresholded GFP signals were indexed as a colour ramp (blue - red for low - high signal intensities). The mitochondria-miniSOG labelling is also color-indexed as dark brown-white for low-high signal intensities. As shown in the first video sequence, other non-PN neurons are also present in the Mz19-GAL4 expression pattern, anterio-dorsal to the AL. To visualise mitochondria-miniSOG localised to PN neurons, a mask (dark purple) was generated from the AL-specific, GFP-positive traced region and applied to the mitochondria-miniSOG image stack. Note that a large fraction of labelled mitochondria aggregated at the cell somata.

**Video 2: Segmentations based on bulk thresholding of mitochondria labelled miniSOG**

This video accompanies Figure 3A-B. The first sequence illustrates the segmented mitochondria (ROI3, cyan-green) obtained from the EM volume, viewed across a 2D stack sequence and then rendered throughout the volume in 3D. In the second clip, a 3D mesh is created from the segmented objects, where individual structures are colour-indexed according to size; blue – green - red; smallest (0.0043 μm^3^) – largest (0.2785 μm^3^). The meshed 3D structure can be compared section-section in 2D against the ultrastructure (inset video). For example, the large red mitochondrion (*1*, in Figure 3B) that appear to be breaking down, as internal cristae no longer exist and miniSOG labels appear aggregated. For comparison, a second smaller mitochondrion is shown, corresponding to *2* in Figure 3B.

**Video 3: Seed-based segmentation of a single cytosol-miniSOG labelled PN cell**

This video accompanies Figure 3C and initially shows a partial stack of the EM volume labelled with cytosol-miniSOG (*1411279R01*, surface 1-180 sections out of the total of 541; volume: 85×85×4.5 micron). These were image-processed by thresholding and Gaussian blur (sigma=3) and two rounds of Sharpen using Fiji. Note the miniSOG contrast on a whole PN cell body and on many labeled neurites that appear contiguous throughout subsections of the AL. From the volume, other partially imaged labeled cell bodies are also present, at the surface and on the right side of the stack. In the segmentation stage, each intended ROI are indicated by the highlighted range values (red). 12 trials were performed, sampling pixels at increasing lower intensities but set to a common maximum value. Note the increasing length and complexity of 3D structures segmented for each ROI.

**Video 4: Further identification of cytosol-miniSOG labelled neurites within volume**

This clip shows some of the other neurite traces found in the image volume, as described in Figure 3D.

**Video 5: XRM image scans of wholemount fly brain**

As described in Figure 4B, some of the cell and neuroanatomical features seen in the XRM scans on a whole fly brain, shown from different planes.

**Video 6: XRM scans of whole fly brain containing cytosol-miniSOG labelled PNs**

This clip shows miniSOG labelled PNs seen in the XRM scans in the fly brain, as described in Figure 4C.

**Video 7: XRM-FIBSEM acquisition workflow**

This clip illustrates the correlative XRM-FIBSEM workflow, which uses non-destructive XRM to screen miniSOG labeled PN neurons in a whole fly brain, followed by FIBSEM volume acquisition, as described in Figure 4.

**Video 8: Examples of over-segmentation errors in cytosol-miniSOG labelled PNs within the AL**

These two video sequences illustrate some examples where it was not possible to fully segment neurites due to signal drop-off. In the first sequence, this occurs as pixel intensities were either very low or had labelling discontinuities along the dendritic segments. This was frequently observed in distal branch segments. In some cases, such discontinuities are considered minor and so merge functions can be easily performed, as illustrated.

The second clip shows images of labelled PN axons in the posterior half of the AL. Here over-segmentation occurs due to the incomplete labelling pattern of cytosol-miniSOG in large axonal segments.

In both cases, having both cell biological and neuroanatomical expertise and the underlying ultrastructure helps to resolve these errors.

